# Integrated microdroplet workflow for high-throughput cell-free transcription in double emulsion picoreactors

**DOI:** 10.1101/2025.11.13.688266

**Authors:** Kartik Totlani, Essa Ahsan Khan, Husnain Ahmed, Rahmi Lale, Bjørn Torger Stokke

**Affiliations:** Biophysics & Medical Technology, Department of Physics, Norwegian University of Science and Technology (NTNU), NO-7491 Trondheim, Norway; Division of Microbial Biotechnology, Department of Biotechnology & Food Science, Norwegian University of Science and Technology (NTNU), NO-7491 Trondheim, Norway

**Keywords:** in vitro transcription, ultrahigh throughput screening, stepinjection, double emulsions, flow cytometry, mango III aptamer

## Abstract

Precise characterization of regulatory sequence performance is fundamental to synthetic biology and next-generation gene therapies, driving the need for scalable and quantitative screening of genetic libraries. While droplet-based microfluidics offers the ultra-high throughput required to scale these assays, it often depends on complex, custom fluorescence-activated droplet sorting platforms. To address this limitation, we introduce an integrated microfluidic workflow that enables cell-free transcription in water-in-oil-in-water double emulsion picoreactors compatible with commercial flow cytometers. The core innovation is an integrated device that combines emulsion reinjection, electric-field-mediated step-injection of in vitro transcription (IVT) reagents, and downstream double emulsification, thereby reducing manual handling and preserving droplet integrity across multistep workflows. We validate the system by coupling isothermal rolling circle amplification (RCA) of DNA templates with on-chip IVT and Mango III aptamer-based fluorescence readout, demonstrating robust detection, binning, and sorting of transcription-active droplets. This workflow provides an accessible and modular platform for quantitative, high-throughput functional screening of regulatory sequences without the need for specialized optical sorting instrumentation.

## 1. Introduction

Single-molecule nucleic acid assays and single-cell gene expression studies have become indispensable for understanding the complexity and heterogeneity of biological systems [1–3]. By analyzing molecular events at the individual molecule or single-cell level, subtle variations that ensemble measurements often obscure can be detected [4]. These high-resolution methods provide critical insights into gene regulation, protein synthesis, and cellular responses to stimuli [5]. Furthermore, the rapid advancements in mRNA therapeutics driven by recent global efforts highlight the need for precise control and characterisation of gene expression [6]. The ability to accurately characterize expression levels, which in turn enables fine-tuning of genetic constructs, is essential for designing optimized systems, ensuring therapeutic efficacy, and minimizing unwanted side effects. [7].

A key factor in regulating gene expression is the strength of the promoter, the *cis*-acting DNA sequence that initiates transcription and determines when, where, and at what level genes are transcribed [8, 9]. Characterising promoter strength is fundamental for designing genetic circuits with predictable expression outputs and remains a central task in synthetic biology [10]. Recently, we described a fluorescence-based bulk assay that uses an RNA aptamer readout to map promoter strength and specificity across large synthetic libraries [11]. While this approach provides quantitative measurements at scale, extending such assays to tens of thousands of regulatory variants requires high-throughput compartmentalisation rather than bulk processing. Microdroplet-based workflows offer this capability: by encapsulating individual transcription reactions into picolitre-scale droplets, each droplet functions as an isolated “bioreactor” containing a single promoter variant. These droplets can subsequently be sorted according to fluorescence intensity, allowing promoter variants to be resolved through quantitative binning rather than broad weak-to-strong assignments.

Droplet-based microfluidics has transformed high-throughput biochemical screening by enabling the generation and manipulation of vast numbers of picolitre to nanolitre compartments [12–14]. It offers ultra-high throughput and the ability to perform complex, multistep protocols in a controlled manner. However, the practical adoption of droplet microfluidics in bioengineering laboratories is often limited by the complexity of the instrumentation. Standard fluorescence-activated droplet sorting (FADS) requires custom-built optical setups, real-time field-programmable gate arrays (FPGAs), and sophisticated alignment, which creates a significant barrier to entry [15]. A promising alternative is the generation of water-in-oil-in-water (W/O/W) double emulsions that are compatible with commercial FACS instruments [16]. While high-performance single-emulsion sorters have been demonstrated, extending these systems to multiplexed or indexed sorting typically requires complicated device fabrication and hardware programming [17, 18]. In contrast, W/O/W double emulsion can be processed on standard benchtop cytometers found in most core facilities, democratizing access to high-throughput screening [19–22].

In this work, we present a modular droplet-based workflow that leverages two separate microfluidic devices to screen in-vitro transcription (IVT) reactions using a fluorescence-based RNA aptamer readout. The schematic overview of the workflow is illustrated in Figure.1 (panels a–f). The workflow comprises: (a) compartmentalization of plasmid DNA in picolitre droplets; (b) off-chip amplification of DNA templates; (c) emulsion reinjection; (d) stepinjection of invitro transcription reagents, (e) conversion of droplets to stable W/O/W double emulsions; and (f) fluorescence-based sorting on a standard FACS instrument. The use of the Mango III RNA aptamer as a transcriptional readout enables direct monitoring of transcript formation and provides a generalisable basis for promoter strength assessment.

**Figure. 1.**
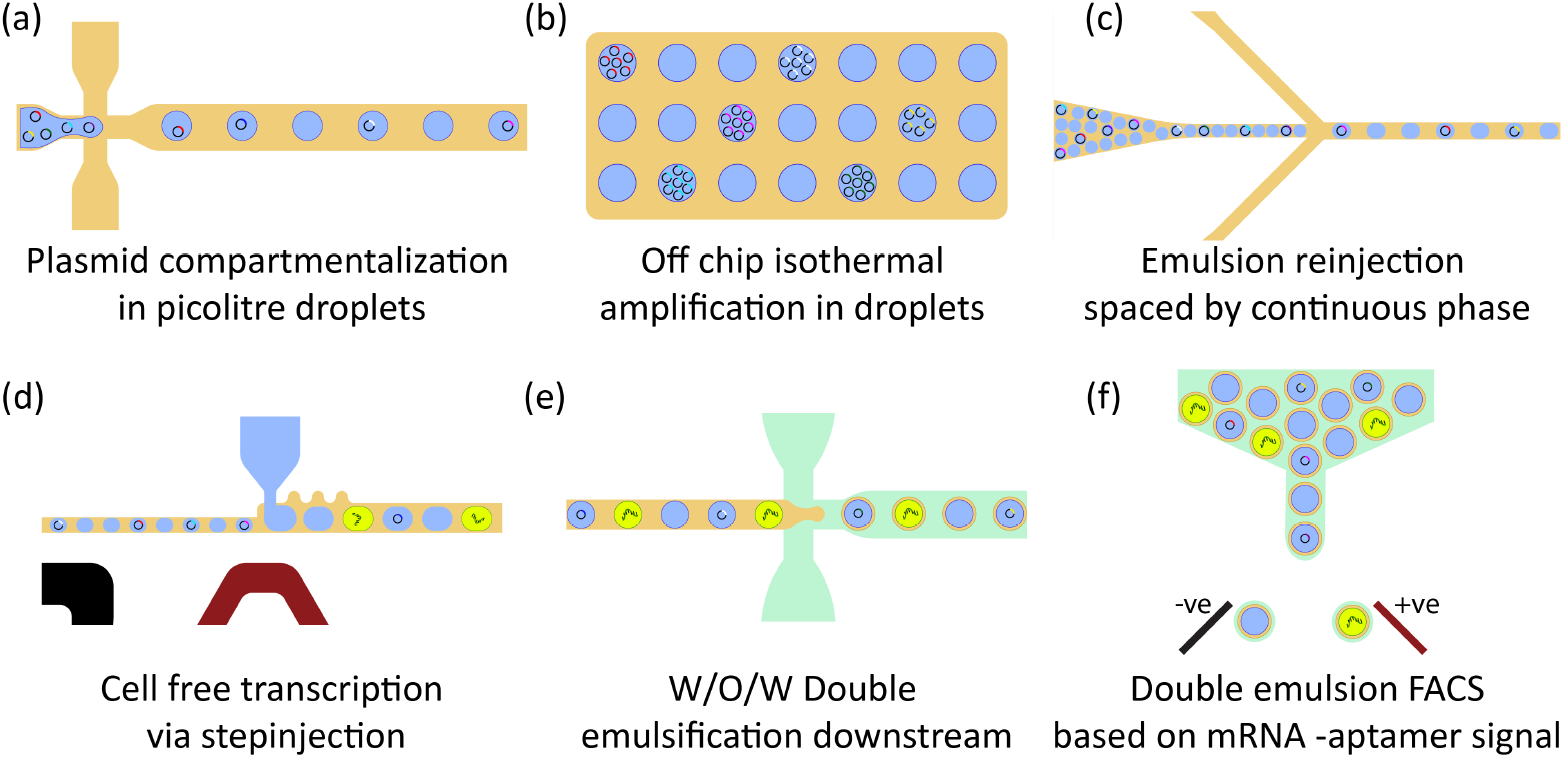
Schematic illustration of the integrated, six-step droplet-based microfluidic workflow for high-throughput screening of nucleic acids using a fluorescence reporter. (a) Plasmid DNA templates are compartmentalised into picolitre droplets. (b) The droplets are incubated off-chip for isothermal rolling circle amplification (RCA) of plasmid DNA. (c) After amplification, the monoclonal droplets are re-injected into a second device and spaced by oil, after which (d) in vitro transcription (IVT) reagents are introduced by electric-field-mediated step-injection to ensure minimal contamination. (e) Each droplet is subsequently converted into a stable water-in-oil-in-water (W/O/W) double emulsion downstream. (f) The double emulsions are then sorted according to fluorescence signal (e.g. Mango-III aptamer readout) on a commercial FACS instrument, enabling intensity-based binning based on the extent of mRNA synthesis.

A key innovation of this platform is the seamless integration of three critical droplet operations: emulsion re-injection, reagent step-injection, and double emulsification onto a single microfluidic chip. While previous work [23] introduced the cross-contamination-free step-injection system, our workflow extends this capability by coupling it directly with on-chip double emulsion formation, thereby minimising manual handling, reducing sample loss, and preserving droplet integrity throughout the multistep protocol. We characterise device performance and validate the workflow using the Mango III RNA aptamer system (as established in [11]) as a transcriptional reporter, demonstrating that transcription-active droplets can be reliably generated, detected, and sorted. As a proof-of-concept, this integrated design establishes a robust and accessible basis for future large-scale, quantitative, and high-throughput characterisation of synthetic regulatory sequences.

## 2. Materials & Methods

### 2.1. Microfluidic device fabrication

#### Device design

The first microfluidic chip, designed for plasmid compartmentalization, featured a simple dual-inlet flow-focusing (FF) junction for droplet generation. The nozzle of the junction was 10 *μ*m wide, and the main channel width was 20 *μ*m. The second microfluidic device incorporated three functional components: (a) emulsion reinjection, (b) a step-injection junction, and (c) a downstream flow-focusing junction for W/O/W double emulsification. Three different microfluidic devices (Design - A, B & C) were designed to test and characterize step-injection and double emulsification (see Figure.3). The channel dimensions for design A and design B are specified in ESI†. Design C was found optimal for the cell free transcription experiments in droplets explored here. Here, the main channel was 13 *μ*m wide upstream of the SI junction and expanded to 25 *μ*m downstream. The SI junction nozzle was 10 *μ*m wide. The corrugated, teeth-shaped structures were integrated adjacent to the nozzle to prevent partial wetting during reagent injection. A serpentine resistor channel was introduced between the stepinjection and double emulsion junctions to electrically isolate the SI junction and prevent corona discharge induced hydrophilicity. The detailed designs and dimensions of both microfluidic devices are provided in Figure.S1 (a & b) of the ESI†.

#### Photolithography

Both microfluidic devices were fabricated using standard photolithography techniques. Briefly, the designs were drafted in AutoCAD 2018 (Autodesk) and converted into direct-writeable DXF files using Layout Editor and Clewin 4.0. The plasmid encapsulation (FF - droplet formation) device was constructed as a single layer microfluidic structure, whereas the second integrated device was constructed in two layers. A negative tone photoresist (mr DWL 5, Micro-Resist technology GmbH) was spin coated on clean 4” silicon wafers as per the manufacturer’s guidelines, followed by soft baking (at 50 and 90°C) to ensure complete solvent evaporation. The structures were patterned via direct laser writing with MLA 150 (Heidelberg instruments) at 405 nm with exposures of 300 - 375 mJ/cm^2^. The post exposure bake was performed at 90°C. To remove any un-exposed photoresist, the resulting microfluidic structures were then developed in mr-DEV 600 (Micro-Resist technology GmbH) and subsequently washed in IPA and dried with nitrogen. Before using the wafers as molds for PDMS devices, their surfaces were silanized by exposing them to vapours of 1H, 1H, 2H, 2H - perfluorooctyltrichlorosilane in a desiccator. The resulting channel heights for the droplet formation devices were measured to be 10.5 *μ*m, while for the step-injection devices, the heights were 10.75 *μ*m for Layer-01 and 10 *μ*m for Layer-02, as determined using a profilometer (Dektak).

#### Soft Lithography

After fabricating the master molds, the PDMS devices were prepared by mixing the elastomer and curing agent (Dow Corning, Sylgard 184 elastomer kit) in the ratio of 10 : 1 (w/w). The mixture was thoroughly mixed, degassed, and poured over the master wafers placed in a 5-inch petri dish. The assembly was cured overnight in an oven at 65°C. Once cured, the PDMS layer was gently peeled off from the wafer and diced. All the inlets and outlets were punched with a 1 mm biopsy punch (Harris Uni-core) and the PDMS molds were thoroughly washed with Isopropyl Alcohol (IPA) and ethanol to get rid of dust and PDMS debris. The resulting molds were then bonded to PDMS spin coated glass slides (25 mm × 75 mm), after exposing both the surfaces to oxygen plasma (Diener Electronic) for 140 s at a pressure of 0.4 mbar. The bonded devices were subsequently baked at 150°C for at least 4 - 8 hours to restore the hydrophobicity of the channels.

#### Microelectrode fabrication

Microelectrodes for the integrated step-injection devices were fabricated by injecting a low-melting-point solder (Indalloy # 19 - 51In, 32Bi, 16.5Sn) into custom-made channels near the step-injection junction. Following device baking, the chips were placed on a preheated hot plate maintained at 95 - 100 °C. Approximately 100 - 200 *μ*L of the molten solder was drawn into a syringe and slowly injected into the electrode channels to ensure bubble-free and uniform filling. Excess molten solder was carefully removed, and the metal connectors (1238060000, male header) were inserted to terminate the electrodes. The devices were thoroughly cleaned, and electrical connectivity was verified using a digital multimeter to ensure reliable and reproducible electrode functionality (see Figure S.2 (a) in the ESI†).

#### Device coatings

Surface wettability was carefully tuned across the two microfluidic devices to ensure optimal performance of each droplet operation. The microfluidic chip used for DNA compartmentalization and droplet formation was rendered hydrophobic to promote stable generation of monodisperse water-in-oil (W/O) droplets. This was achieved by filling the channels with 1 % (v/v) 1H, 1H, 2H, 2H - Perfluorooctyl - trichlorosilane dissolved in fluorinated oil (Picowave 7500, Sphere Fluidics). Excess solution was removed using compressed air and the devices were baked at 150°C prior to use.

In contrast, the second device which integrates emulsion reinjection, step-injection (SI), and downstream W/O/W double emulsification required spatially selective surface modifications. Firstly, the SI junction and the emulsion reinjection region were coated hydrophobic using the method described above. To enable reliable oil phase pinch-off and robust double emulsion formation, the right-hand side of the W/O/W - FF junction was rendered hydrophilic. This asymmetry in surface wettability was essential for facilitating water-in-oil-in-water (W/O/W) droplet formation and preventing wetting failures. The hydrophilic surface functionalization was achieved in two steps. First, the junction was exposed to a locally generated corona discharge [24] by inserting the tip of the electrode through the channel outlet and activating the electrode for 10 seconds. Next, the junction was chemically modified using a sequential layer-by-layer coating with polyelectrolyte solutions to enhance hydrophilicity [25]. Briefly, after the corona discharged exposure, the channel was flushed with 2 mg/mL solution of PDADMAC (poly (diallyldimethylammonium chloride)) prepared in 0.5M NaCl for 10 minutes followed by rinsing with 0.1 M NaCl. Subsequently, the channel was flushed with 2 mg/mL solution of PSS (poly(styrene sulfonate)) in 0.1 M NaCl for 10 minutes and finally, the channel was flushed with filtered MilliQ water. The entire procedure for preferential coating of step-injection devices has been illustrated in Figure S.2 (b) in the ESI†.

### 2.2. Working fluids

#### Continuous phase

Fluorinated oil was used as the continuous organic phase for all the experiments. For droplet formation and DNA compartmentalization, 3.5 % Picosurf-1 (Spherefluidics) dissolved in fluorinated oil Picowave - 7500 (Spherefluidics) was used as the continuous phase. For spacing the emulsion during the re-injection and reagent step-injection, 1% (v/v) Picosurf-1 was used as the continuous phase. For the formation of Water-in-Oil-in-Water (W/O/W) double emulsion droplets downstream, a 2 % Tween-80 solution in MilliQ water was used as the continuous aqueous phase. All the continuous phase solutions were filtered through 0.2 *μ*m pore size filters before injection.

#### T-7 Mango plasmid construction

The Mango III aptamer, synthesized by Twist Bioscience, was first amplified using the Lib1a-F and Lib2a-R primers. The T7 promoter was then incorporated using the Lib1b-F and Lib2b-R primers. The resulting T7-Mango III product was further amplified with the Lib3-F and Lib4-R primers for Gibson assembly into the pSEVA2311 backbone, which had been previously amplified using the Lib5-F and Lib6-R primers. The assembly was performed using the NEBuilder HiFi DNA Assembly Master Mix. Successful integration of the T7-Mango III-pSEVA2311 construct was confirmed through colony PCR with the Lib7-F and Lib8-R primers before being used for the IVT reaction. The primers has been listed in Table.1 of the ESI†.

#### Isothermal RCA mix

For isothermal amplification in droplets, the illustra GenomiPhi V2 DNA Amplification Kit (Cytiva) was utilized. This kit provides the necessary buffers, random hexamers, and Phi29 DNA polymerase required for plasmid amplification. Briefly, the desired concentration of T7-Mango III-pSEVA2311 template was mixed with the sample buffer in a 1:9 (v/v) ratio. The mixture was heated at 95°C for 3 minutes and then immediately cooled on ice. Separately, the Phi29 DNA polymerase was combined with the reaction buffer in another tube also in the ratio of 1:9 (v/v). Both tubes were cooled on ice before being introduced into the microfluidic device for droplet emulsification.

#### In-vitro transcription in droplets

The MAXIscript™ T7 Transcription Kit (Invitrogen, Thermo Fisher) was used to prepare the in vitro transcription (IVT) reagent mix. The mix consists of nuclease-free water, 10X transcription buffer, 10 mM of each rNTPs, KCl, and T7 RNA polymerase. To visualize the mRNA transcripts generated within the droplets, TO1-3PEG-Biotin fluorophore (Applied Biological Materials) was included in the reagent mix. This dye specifically binds to the Mango aptamer incorporated into the mRNA, producing a strong fluorescence signal (Excitation wavelength of 505 nm and emission wavelength of 540 nm). A concentration of 150 mM KCl was required to facilitate the binding of the Mango aptamer to the Biotin-conjugated fluorophore, enabling sensitive detection of mRNA transcripts within the droplets. For droplet-based experiments, the IVT stoichiometry was adjusted to account for StepInjection dilution and to maintain a final 1X buffer strength inside the picodroplets. Accordingly, a 120 *μ*L IVT master mix was prepared using the following component volumes: 44 *μ*L nuclease-free water, 16 *μ*L of 10X transcription buffer, 16 *μ*L of 150 mM KCl, 4 *μ*L TO-1 Biotin (1:16.5 stock), and 8 *μ*L each of ATP, CTP, GTP, UTP (10 mM stocks), along with 8 *μ*L T7 polymerase enzyme. This modified formulation ensured that, after injection, the reagent composition inside the droplet matched the required working concentrations without exceeding ionic strength limits or compromising droplet stability. If the injection volume or droplet size is changed, the IVT formulation must be recalculated proportionally to maintain the correct final 1X buffer condition and the required concentrations of rNTPs, KCl, and buffer components inside each droplet.

### 2.3. Microfluidic operations, image acquisition & analysis

The fluids were injected using the Nemesys S (Cetoni) and PHD Ultra (Harvard Apparatus) syringe pumps. Gastight luer lock glass syringes (1 mL : 1001 TLL and 5mL : 1005 TLL, Hamilton) were used to drive the fluids. Medical grade PTFE tubing (ID : 0.6 mm, OD : 1.07 mm, 24G, BB311-24, Scientific commodities inc.) was used to connect the syringes to the microfluidic chips via precision dispensing needle connectors (ID : 0.33 mm, 23G, L : 6.4 mm, 923025, TE Metcal). For injecting small volumes of the aqueous phases (e.g. 50 - 150 *μ*L), the solutions were withdrawn into a pre-filled tubing and syringe with an air bubble gap to maintain separation. For the stepinjection, on-chip electrodes were connected to a high voltage amplifier (623B, TREK) via copper wires connected to a metal connector (1281760000, female header, Weidmüller). The high voltage amplifier amplified the signal from a waveform signal generator (33600A, Keysight Technologies) before transmitting them to the on-chip electrodes. The AC electric potential was maintained at a frequency of 30 kHz and an amplitude of 120 *V*_*PP*_ to ensure a reliable injection of IVT reagents into the passing droplets. A high-speed camera (FASTCAM SA3, Photron, model: 120 K M1) mounted on an inverted microscope (Nikon Eclipse Ti2) was used to observe the droplet microfluidic experiments and record high-speed movies. Real time validation of droplet generation frequency and volume during the experiments was performed by recording at 30,000 FPS. The edge detection analysis for the droplet volume measurement was performed by Droplet Morphometry and Velocimetry (DMV) [26]. The droplet volumes before and after stepinjection were calculated using the 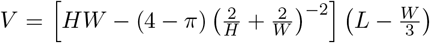 [27], where the L represents the length of droplet along the major axis as detected by DMV and H, W are the height and width of the microfluidic channel respectively. Fluorescence and brightfield images of double emulsion droplets both before and after FACS sorting were acquired using a Zeiss LSM 800 Airyscan confocal microscope, allowing simultaneous visualization of droplet morphology and the FITC-channel fluorescence arising from TO1-3PEG-Biotin Fluorophore-Mango complex generated during IVT. Imaging was performed using the 495 nm excitation / 519 nm emission FITC channel on the LSM 800 URGB laser module (10 mW diode laser), with signal collected on internal GaAsP photomultiplier detector. Depending on the required field of view and resolution, either a 10X air objective (C-Apochromat 10X/0.45) or a 20X water-immersion objective (Plan-Apochromat 20X/0.8) was used.

### 2.4. Double emulsion sorting on FACS

The W/O/W double emulsions post IVT incubation were analyzed and sorted on the SH800S Cell Sorter (Sony Biotechnology). Before the injection, a 50 *μ*L DE emulsion was diluted in 500 *μ*L of 1X sheath buffer (Sony) in a 12 × 75 round bottom FACS tube. The emulsion was interrogated using a standard blue laser (488 nm) configuration and a 130 *μ*m sorting chip. This sorting nozzle size leads to approximately 8 nL drop post pinch off with the effective sorting frequency of around 12 kHz. FSC was used as the trigger with a detection threshold of 0.67 %. The FSC gain was 1, BSC gain 28 % and the gain for Mango signal was 40 %. The sample injection pressure was kept high (8 - 9 psi) in the initial phase. After a brief pickup delay, double emulsions appeared as the events and were gated on BSC-A vs. FSC-A for gating and analysis. Once the events appeared the pressure was reduced to 4 - 5 psi. The event rate was kept between 300 - 800 events/s to minimize any shear induced breakup of the double emulsions. Since the DEs were denser than the sheath buffer, they were manually suspended intermittently during the analysis and sorting. The double emulsion initially gated on the BSC vs FSC signals, were subsequently gated on Mango-TO1-3PEG biotin signal on the fluorescence histogram.

## 3. Results & discussions

### 3.1. Double emulsion pico-reactor workflow

#### Droplet formation and encapsulation of T7-Mango plasmids

The droplet-based screening workflow begins with co-encapsulation of T7-Mango plasmids and the isothermal RCA regents into picolitre-sized droplets using a dual-inlet flow focusing microfluidic device. Plasmids and sample buffer were injected via the first inlet, while enzyme and reaction buffer were introduced via the second (see Figure.S1 (a) of the ESI†). Figure. 2 (a) shows the droplet formation process. The startup protocol involved prefilling the microfluidic device with the continuous oil phase at a flowrate of 100 - 200 *μ*L/h. Once the stable oil flow was established, both the aqueous phases were injected at 25 *μ*L/h ensuring that dispersed phases did not wet the PDMS channel walls. This approach consistently enabled stable emulsion production with uniform droplet volumes and no partial wetting. Following the initialization, the steady state flowrates were optimized to 75 *μ*L/h for the continuous oil phase and 25 *μ*L/h for both the aqueous phases (50 *μ*L/h in total). Under these settings, the device reliably produced droplets of ∼ 2.3 pL at a generation frequency of ∼ 6.5 - 7 kHz in the *dripping* regime ensuring a high droplet monodispersity (*<*1 % COV). While the frequency and droplet volume during the experiment was confirmed by recording high speed movies, the movies were also analyzed via DMV for droplet volume calculations via edge detection [26] (see Figure. 2 (b)). The droplet size distribution shown in Figure.2 (d) exhibits a narrow unimodal peak centered at 19.9 ± 0.05 *μ*m, with a coefficient of variation below 0.5 %, confirming the high monodispersity of the emulsion. The corresponding droplet volume within the channel geometry is estimated to be 2.3 pL, consistent with the flowrate-controlled breakup mechanism governing the dripping regime in confined microfluidic geometries, as established by Garstecki et.al. [28].

**Figure. 2.**
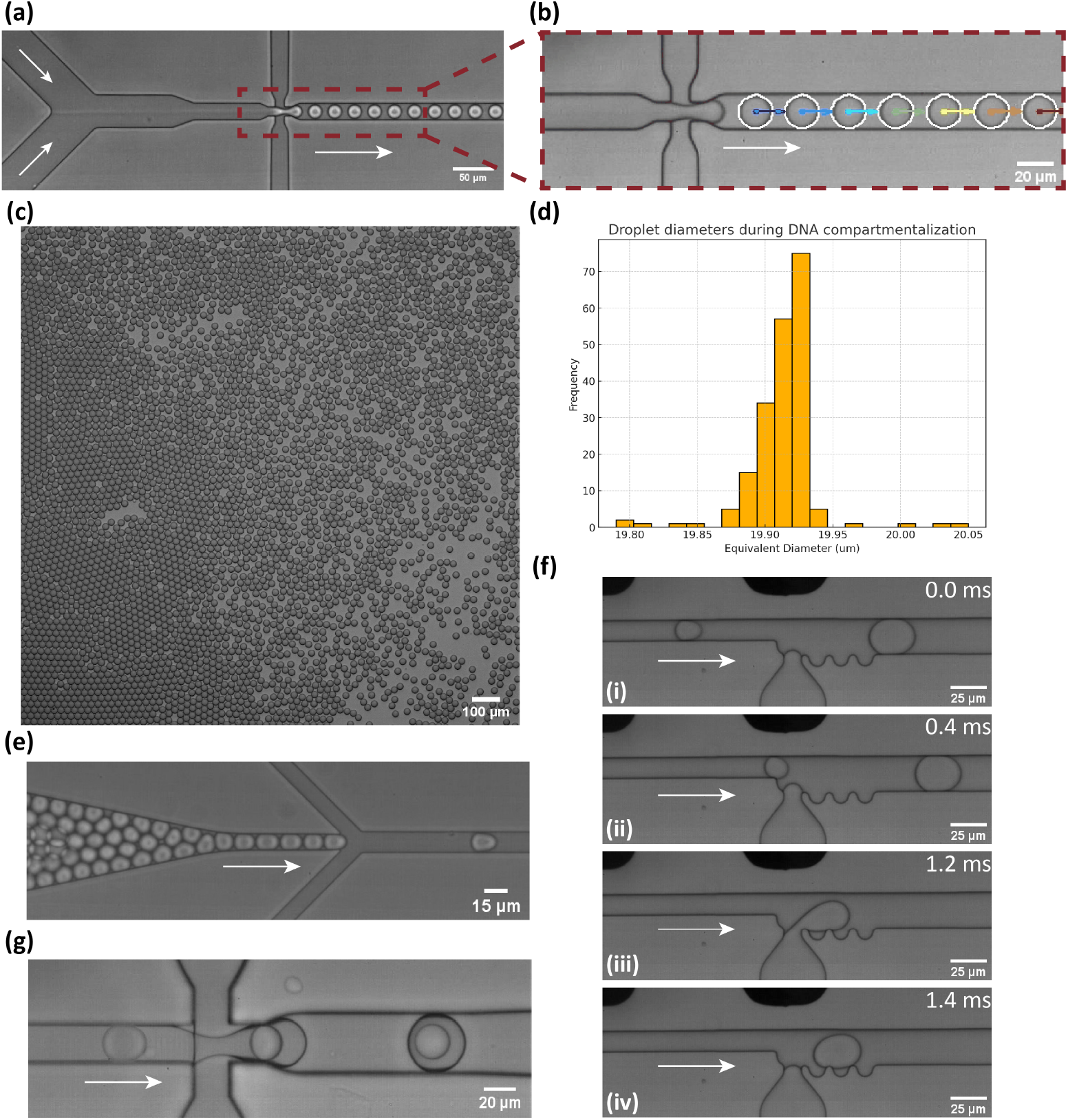
Integrated microfluidic workflow for high-throughput cell-free transcription in double emulsion picoreactors. (a) Droplet generation using a dual-inlet flow-focusing junction for encapsulating T7-Mango plasmids and RCA reagents. (b) A video frame used for droplet size and monodispersity quantification via edge detection in the DMV analysis [26]. (c) A micrograph of emulsion after isothermal rolling circle amplification (RCA), confirming surfactant stability and droplet integrity. (d) Histogram showing droplet size distribution after compartmentalization and RCA (n = 200 droplets), confirming *<*1 % COV. (e) Reinjection of droplets into the integrated chip, spaced by 1 % Picosurf-1 in HFE-7500. (f) Time-lapse series of electric field-mediated StepInjection of IVT reagents (T7 RNA polymerase, rNTPs, and TO1-3PEG-Biotin Fluorophore) into flowing DNA-containing droplets. (g) Downstream W/O/W double emulsification of injected droplets using a 2 % Tween-80 continuous aqueous phase, yielding stable double emulsion (DE) picoreactors suitable for off-chip incubation and FACS-based phenotyping.

**Figure. 3.**
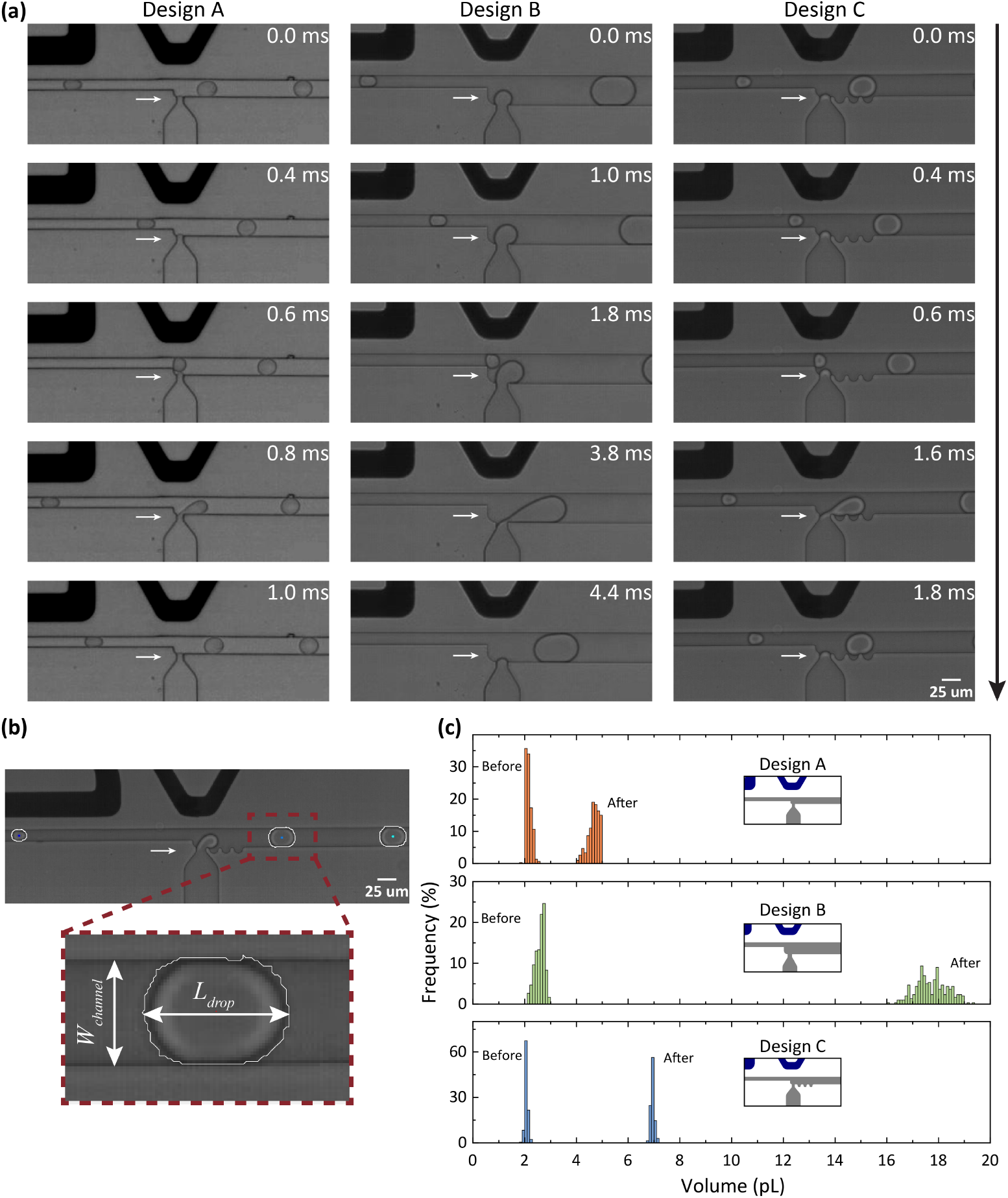
Characterization of microfluidic reagent stepinjection for droplet based in vitro transcription assays. (a) Time-lapse microscopic images showing electric field-mediated stepinjection in three microfluidic devices (Design A, B, and C), where the post-junction channel width and nozzle dictates the injected reagent volume. (b) Representative frame from Droplet Morphometry and Velocimetry (DMV) analysis showing edge detection and droplet length measurement before and after injection. (c) Quantitative assessment of droplet volumes before and after StepInjection across 300 droplets per design. Volume consistency and monodispersity were maintained post-injection.

#### Rolling circle amplification in picodroplets

Emulsion containing T7-Mango DNA plasmid templates and the RCA mix was collected in a 1 mL syringe (B-Braun) prefilled with 3.5 % Picosurf-1 in Picowave-7500. After approximately 2 hours of collection, the droplets were incubated at 30 °C overnight (16 h) to enable isothermal amplification. The Phi29 DNA polymerase in the mix employs random hexamers to initiate rolling circle amplification, generating long concatemeric DNA strands from the circular T7-Mango plasmid template [29, 30]. The enzyme was deactivated by incubating the emulsion at 65 °C for 10 minutes after the amplification.

Compared to conventional thermocycling methods such as ddPCR, RCA offers a gentle and isothermal protocol that preserves droplet stability and integrity [31]. Initial attempts using ddPCR thermocycling with Q5 and TAQ polymerases led to droplet coalescence and loss of monodispersity, likely due to surfactant instability at elevated temperatures (≥ 90°C). This instability was consistently observed across multiple surfactant formulations and temperature profiles (see Figure S.3 in the ESI†), highlights the limitations of thermocycling for droplet-based PCR. The monodispersity of the emulsion remained intact after amplification conducted at lower incubation temperature. Figure.2 (c) shows a highly monodisperse emulsion following amplification which is critical for ensuring a stable emulsion reinjection and further downstream droplet operations.

#### Cell free transcription via stepinjection and double emulsification

Efficient and reliable injection of IVT reagents into DNA-containing droplets is central to the robustness of any droplet-based promoter discovery routine. Typically, in droplet microfluidics, pico-injection [32] is employed for this purpose; however, pico-injectors may be susceptible to inter-droplet cross contamination and can be suboptimal for molecular biology assays that demand high specificity and sensitivity [33]. To address this, we implemented “StepInjection” previously introduced by Hu & co-workers [23]. In this work, we extend this concept by integrating it upstream of a W/O/W double emulsification module, enabling downstream compatibility with standard FACS-based double emulsion phenotyping. Figure S1 (b) in the ESI† shows the complete layout of this integrated device with all relevant channel dimensions. This integration facilitates three operations to be performed on a single microfluidic chip: (i) emulsion re-injection (see Figure.2 (e)), (ii) StepInjection of the IVT reagents (see Figure.2 (f)) and (iii) double emulsification of droplets post StepInjection (see Figure.2 (g)).

The integrated device coordinates four fluid streams to execute the overall multi-step workflow. Firstly, the single emulsion droplets post-isothermal amplification was reinjected into the chip and spaced using 1 % Picosurf-1 in Picowave 7500 forming an emulsion train with controlled droplet spacing (Figure.2(e)). High droplet monodispersity was critical for the stability of subsequent emulsion reinjection. At the step-injection (SI) junction, IVT reagents including T7 RNA polymerase, rNTPs, and the fluorogenic TO1-Biotin dye were injected into each droplet via electric field mediated interface de-stabilization. Figure.2(f) shows a timelapse series of the stepinjection process. The electric potential (30kHz, 120 *V*_*PP*_) was applied through on-chip microelectrodes to drive a contamination free mering of IVT reagents. Immediately downstream, the reagent-injected droplets entered a flow-focusing (FF) junction where a 2 % Tween-80 aqueous stream sheared the oil phase to form stable water-in-oil-in-water (W/O/W) double emulsion picoreactors. This is shown in Figure.2(g). The resulting DEs comprising of an oil shell and a post-injection core were collected and incubated at 37 °C prior to FACS based sorting.

Successful execution of this integrated workflow relied on a strict and deliberate startup protocol to prevent backflow and partial wetting. The downstream FF junction was first primed with the continuous aqueous phase (2 % Tween-80 in MilliQ). Once filled, the spacer oil flow was initiated, producing stable oil-in-water droplets to establish the baseline flow. Only after both continuous phases were stabilized was the IVT reagent stream introduced at the SI junction. At this stage, transient droplet formation at the SI junction was observed until the emulsion re-injection was started. Once the emulsion enters, all four flowrates re-injection, spacer oil, IVT reagents, and continuous aqueous phase were fine-tuned to reach a stable operational regime. The steady state flowrates were *Q*_*em*_ = 6 *μ*L/h for the emulsion, *Q*_*sp*_ = 65 *μ*L/h for the spacer oil, *Q*_*in*_ = 10-15 *μ*L/h for the IVT reagents and *Q*_*aq*_ = 150 *μ*L/h for the continuous aqueous phase(2 % Tween). In this final steady-state, a continuous train of monodisperse single emulsions entered the chip, received precise reagent addition, and were converted into W/O/W double emulsions at the FF junction.

Representative movies illustrating the complete integrated workflow including droplet generation, emulsion reinjection, StepInjection of IVT reagents, and downstream W/O/W double emulsification are illustrated in Movie-1 of ESI†.

### 3.2. Characterizing reagent stepinjection

To evaluate how StepInjection (SI) junction geometry influences reagent injection, we tested three microfluidic designs that varied in the width of the main channel upstream and down-stream of the SI junction as well as nozzle width. All DNA-containing droplets originated from the same upstream generation protocol, ensuring consistent initial droplet volumes across the three designs. Time-lapse microscopy of the SI region for each design is shown in Figure.3 (a), with corresponding channel dimensions annotated. Representative high-speed recordings illustrating the injection dynamics for the different SI geometries are provided in Movie 2 of ESI†. Quantitative volume analysis was performed using edge detection via Droplet Morphometry and Velocimetry (DMV) on high-speed recordings (see Figure.3 (b). For each design, droplet volumes before and after reagent addition were extracted, and the injected volume was calculated as their difference. As shown in Figure.3 (c), the injected volume increased systematically from Design A to Design C to Design B, correlating with the progressive widening of the post-injection channel. This trend aligns with predictions from droplet formation physics in the squeezing regime at low capillary numbers, where a wider downstream channel prolongs the coalescence and pinch-off time, thereby permitting a larger volume of reagent to be incorporated before detachment occurs [34, 35]. While Design A enabled high-frequency injections, the limited reagent volume (∼ 6 pL) may be insufficient for robust IVT, increasing the risk of false-negative readouts. Conversely, Design B achieved the largest injection volumes (∼ 18 pL) but exhibited slower droplet processing and instability during W/O/W double emulsification, likely due to larger internal droplet volumes challenging the pinch-off at the final flow-focusing junction. Design C, with an intermediate channel geometry, achieved a favorable tradeoff: sufficient reagent addition for reliable IVT, stable double emulsion formation, and throughput compatible with large-scale screening. Importantly, all three design variations delivered tight monodispersity of droplets after injection, confirming the consistency of SI performance across geometries. These findings emphasize the critical role of channel width following the junction site in tuning reagent delivery and throughput, allowing the SI architecture to be tailored for assay-specific demands.

### 3.3. Double emulsion phenotyping via FACS

To phenotype and sort droplets based on IVT-driven fluorescence, water-in-oil-in-water (W/O/W) double emulsions were analyzed using a commercial FACS system (Sony SH800) equipped with a 488 nm excitation laser and FITC detection channel. The overall gating strategy is summarized in Figure.4. First, DE droplets were distinguished from oil droplets, debris, and background based on their characteristic forward scatter (FSC) and side scatter (SSC) signatures (see Figure.4 (a)), reflecting their larger size and composite core-shell structure. Next, singlet DE droplets were isolated by gating on FSC height versus FSC width (see Figure.4 (b)), thereby excluding coincident events and droplet aggregates that could confound fluorescence quantification. Following singlet selection, fluorescence intensity distributions in the FITC channel were used to identify droplets exhibiting Mango aptamer signal arising from successful IVT (see Figure.4 (c)). A threshold gate was defined based on negative-control populations to separate fluorescent-positive and fluorescent-negative DE droplets. Finally, gated populations were visualized in FSC-FITC space (see Figure.4 (d)) to verify consistent separation prior to sorting. This multistep gating approach, similar to previously reported DE-FACS workflows [20, 21], enables reliable discrimination of intact double emulsions, removal of multiplets, and quantitative assessment of IVT activity at the single-droplet level, forming the basis for downstream fluorescence-based sorting and indexing.

**Figure. 4.**
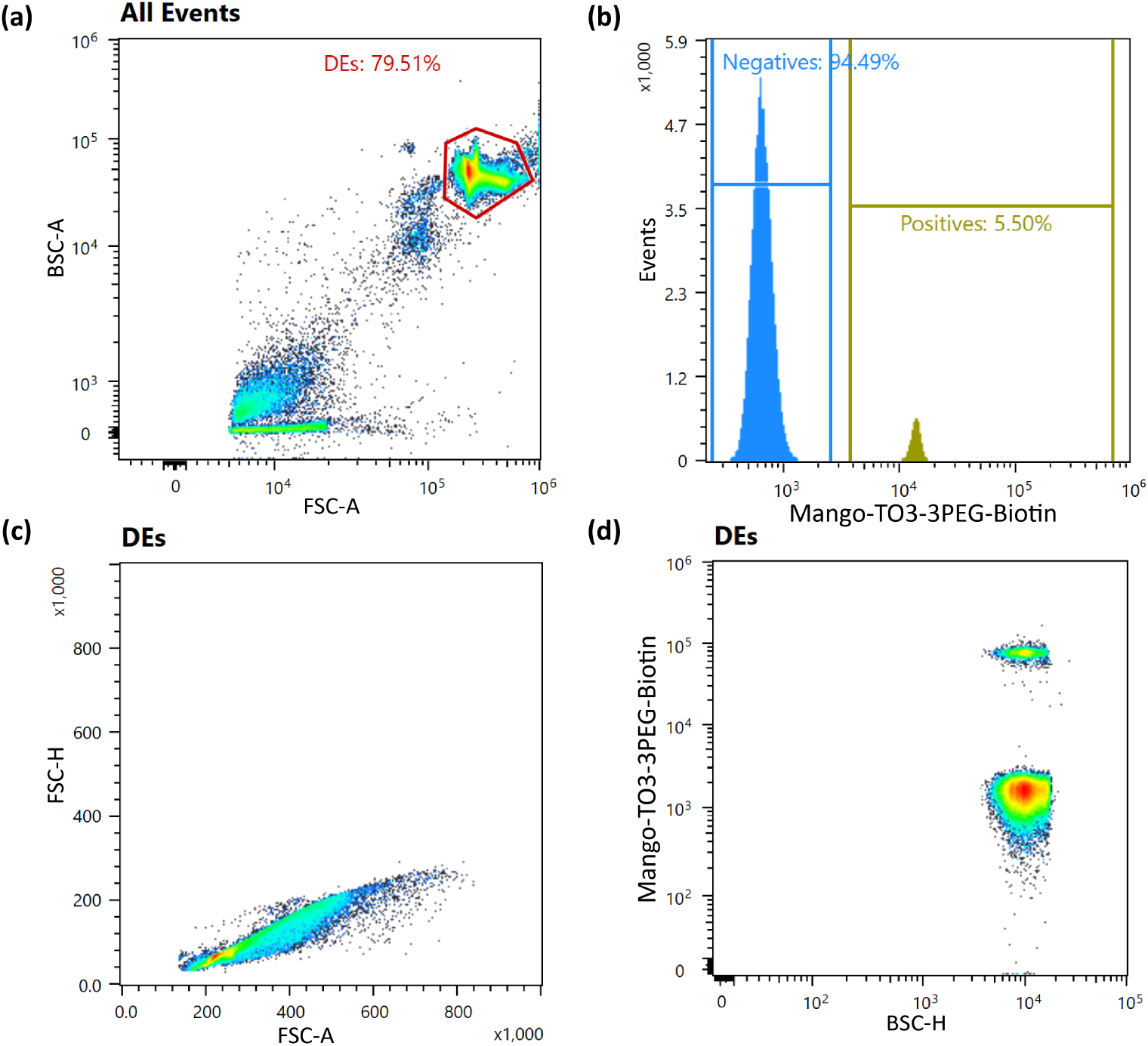
(a) Back scatter (BSC-A) vs. forward scatter (FSC-A) plot of all detected events. The gated population (red polygon) corresponds to DE droplets based on their expected size and granularity, comprising 79.51 % of total events. Smaller oil droplets and debris appear as lower FSC-A / BSC-A populations and are excluded. (b) Fluorescence histogram (FITC-H channel) of gated DE droplets showing distinct negative (blue) and positive (olive) populations. Linear gating separates the fluorescently active DEs (5.5 %) from the non-fluorescent background (94.5 %) for downstream sorting and analysis. (c) Doublet discrimination plot of FSC-H vs. FSC-A for DE droplets. The diagonal trend indicates a predominantly singlet population with minimal doublet contamination. No additional gating was applied, as the population lies tightly along the diagonal. (d) Two-dimensional dot plot of FITC-A (fluorescence) vs. BSC-H, showing two distinct droplet populations. The lower cluster represents non-fluorescent DEs, while the upper cluster corresponds to highly fluorescent DEs, corroborating the histogram-based gating.

To validate the functional performance of the integrated DE droplet workflow, we performed a set of proof-of-concept experiments using a known positive control plasmid encoding the T7-Mango aptamer. In this approach, we systematically varied the initial plasmid concentration across three separate experiments, simulating different gene loading scenarios. These dilutions : 1:1000 (low), 1:100 (medium), and 1:10 (high) were applied during the initial DNA encapsulation step. Due to the stochastic nature of encapsulation, droplet occupancy follows a Poisson distribution, resulting in a large fraction of empty droplets, especially at lower input concentrations. Following isothermal RCA, step-injection of IVT reagents, and overnight incubation at 37 °C, the resulting W/O/W DE droplets were analyzed via FACS (Sony SH800) using the FITC channel to detect Mango-TO1 Biotin fluorescence. Figure.5 (a) summarizes the two experimental cases, while Figure.5 (b) shows representative brightfield and fluorescence microscopy images of the resulting DE droplets, where the number of fluorescent cores increases with input DNA concentration. Consistent with this trend, the corresponding FACS histograms in Figure.5 (c) show that the fraction of fluorescent-positive droplets increased with input DNA concentration, from 0.81 % at 1:1000 to 18.59 % at 1:10. These values matched expectations based on Poisson statistics and confirmed that only DNA-loaded droplets exhibited Mango signal following successful transcription. To assess the ability of the platform to isolate functional droplets, we sorted the positive population based on Mango fluorescence and re-imaged the sorted DEs under fluorescence and brightfield microscopy. Figure.5 (d) demonstrates a high level of sorting fidelity: sorted positives consistently showed bright fluorescent cores, while the negative population exhibited no detectable fluorescence. Together, these experiments establish that our DE-based droplet platform enables robust, high-resolution phenotyping of IVT activity at the single-droplet level. Although these experiments involve a known positive control, they validate the key capabilities needed for future high-throughput screening of randomized promoter libraries which are reliable reagent delivery, signal-specific readouts, and efficient recovery of functional variants based on fluorescence signals.

**Figure. 5.**
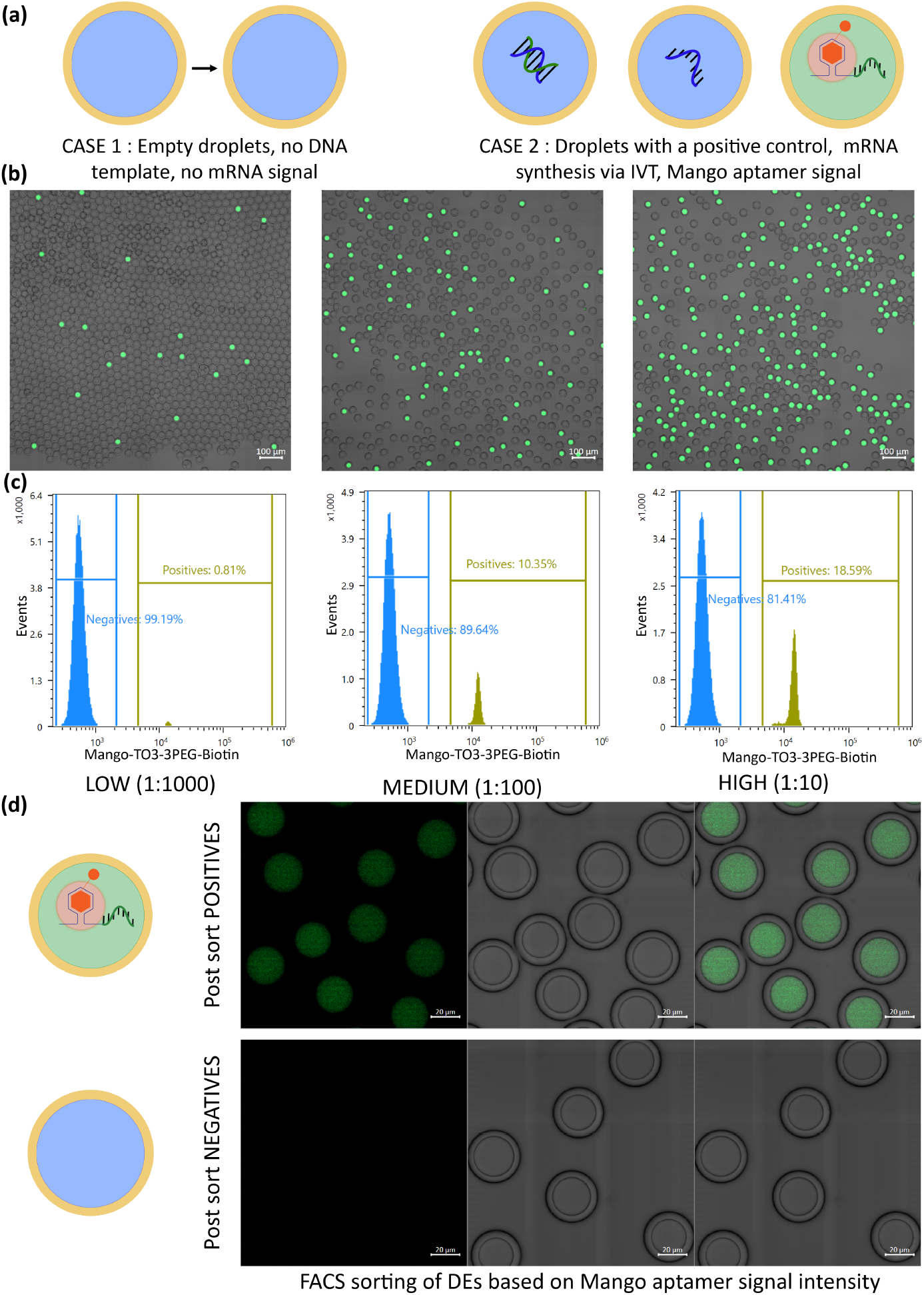
Phenotyping and FACS sorting of double emulsion (DE) droplets based on IVT-driven Mango aptamer fluorescence. (a) Top row: Schematic illustrations of experimental cases : Case 1 (left) shows empty droplets lacking DNA template, while Cases 2 (middle, right) represent droplets encapsulating T7-Mango plasmids undergoing transcription reaction inside the droplet. (b) Overlaid fluorescence and brightfield microscopy images of DE droplets after incubation for the IVT reaction reveal an increase in the number of fluorescent cores with higher input DNA. (c) Corresponding FACS histograms showing clear separation between fluorescent-positive and negative droplets across three template loading levels : low (1:1000), medium (1:100), and high (1:10). As expected from Poisson statistics, the percentage of positive droplets increases with template concentration: 0.81 %, 10.35 %, and 18.59 %, respectively. (d) Analysis of FACS sorter DE droplets based on Mango signal intensity. Top: Positively sorted droplets exhibit bright fluorescent cores, indicating successful transcription reaction.; Bottom: Negatively sorted droplets show no signal, confirming accurate phenotypic gating and separation.

### 3.4. Intensity based droplet binning

Building on this paradigm, we demonstrate a fluorescence intensity based indexed sorting strategy for double emulsion (DE) picoreactors using a commercial fluorescence activated cell sorter (FACS). This workflow parallels the multiplexed droplet sorting concept reported by Caen and co-workers [17] but extends it to water-in-oil-in-water (W/O/W) emulsion offering greater stability, and full compatibility with standard FACS instrumentation. Instead of performing on-chip single emulsion (O/W) microfluidic sorting, the DE droplets are electronically gated and deposited into predefined wells of a Microtiter plate according to their fluorescence intensity levels, enabling accurate binning of transcriptional activity.

As a proof of principle, we used T7-Mango plasmids as a positive control template to generate fluorescence upon successful in vitro transcription (IVT) inside the droplets using the integrated workflow. The resulting Mango mRNA fluorophore complexes produced signals in the FITC channel. As shown in Figure.6 (a), DE droplets were categorized into three fluorescence intensity bins : high, medium, and low; each representing distinct mRNA yield distributions. These categories were indexed to specific regions of a 96-well plate (Figure.6 (b)), and post-sort microscopic imaging confirmed accurate fluorescence-based segregation (Figure.6 (c)). Wells corresponding to high-intensity gates exhibited densely fluorescent droplets, while medium and low bins contained progressively dimmer populations, validating the intensity-based categorization. This experiment establishes the foundation for functional screening of synthetic promoter libraries in which droplets would encapsulate a unique promoter variant driving transcription from a *σ* factor specific RNA polymerase complex. The resulting fluorescence intensities could thus report on promoter strength, allowing rapid categorization of artificial constructs into “strong,” “medium,” and “weak” regulatory elements. By operating under Poisson loading conditions (∼ 20 % occupancy with single variant), this approach would enable single template encapsulation and minimize crosstalk, facilitating genotype-phenotype mapping. This fluorescence-indexed DE sorting framework conceptually complements the bulk in vitro transcription-based promoter characterization we reported previously [11], and illustrates how similar promoter activity measurements could, in the future, be adapted to a single-template droplet format to enable higher-throughput, finer-resolution functional screening.

**Figure. 6.**
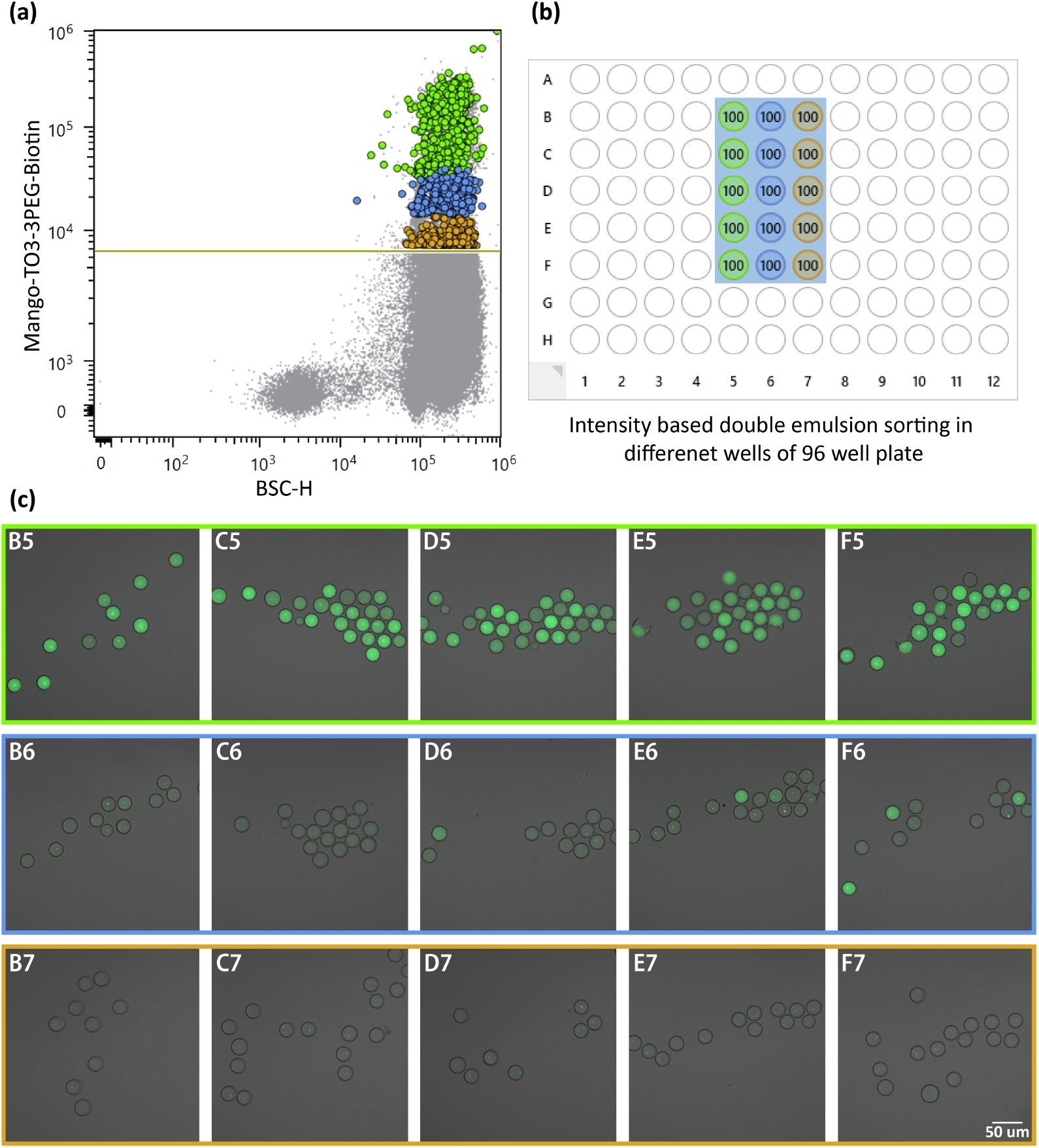
Fluorescence intensity-based indexing of double emulsion (DE) droplets using FACS. (a) Fluorescence gating strategy used to partition DE droplets into discrete intensity bins (high, medium, low) based on Mango RNA fluorescence in the FITC channel. (b) Indexed deposition map showing the predefined well positions corresponding to each fluorescence gate during single-droplet sorting into a 96-well plate. (c) Representative post-sort microscopy images of W/O/W droplets retrieved from individual wells across the three fluorescence categories, illustrating clear separation of bright, intermediate, and dim droplet populations. The figure highlights the ability of the DE workflow to generate distinct transcriptional phenotypes and to recover rare high-intensity events through fluorescence-based binning.

## 4. Conclusions

In this work, we have established an integrated microdroplet workflow for high-throughput cell-free transcription that effectively bridges the gap between complex microfluidic operations and accessible flow cytometry-based phenotyping. By consolidating emulsion reinjection, electric-field-mediated StepInjection, and water-in-oil-in-water (W/O/W) double emulsification onto a single integrated device, we significantly reduced manual handling and minimized sample perturbations. We further characterized the StepInjection dynamics, demonstrating that the post-junction channel geometry is a critical design parameter for tuning reagent delivery volumes.

Validation of the platform using the T7-Mango aptamer system confirmed that transcriptional activity could be initiated, maintained, and detected within the double emulsion picoreactors. The resulting fluorescence distributions aligned with Poisson statistics, validating the platform’s sensitivity and its ability to distinguish positive “bioreactors” from empty droplets across varying template concentrations. Crucially, by generating stable W/O/W double emulsions, this workflow circumvents the need for specialized, custom-built fluorescence-activated droplet sorting (FADS) setups. Instead, it enables the use of standard commercial FACS instruments for the intensity-based binning and recovery of transcriptional phenotypes.

Looking forward, this platform provides the necessary technological foundation to scale functional promoter characterization from bulk assays to high-throughput single-molecule screens. The workflow is immediately applicable to the screening of randomized regulatory libraries, where droplets containing unique promoter variants can be challenged with specific RNA polymerase - sigma factor complexes. With fluorescence intensity serving as a quantitative proxy for transcriptional strength, this system allows for the precise ranking and sorting of genetic elements. Overall, this work establishes a robust, modular framework for genotype-phenotype coupling, unlocking new possibilities for quantitative synthetic biology and regulatory genomics.

## Supporting information

Electronic Supplementary Information (ESI)

Movies and Design files

## Conflict of Interest

The authors declare no conflicts of interest.

## Acknowledgements

We acknowledge funding from the Research Council of Norway (RCN) through the FRIPRO grant (grant No. 316129). We thank the Norwegian Infrastructure for Micro- and Nanofabrication (NORFAB, grant No. 295864) at NTNU NanoLab for providing access to cleanroom infrastructure and microfabrication support. The authors also extend special thanks to Dr. Liisa van Vliet and Prof. Florian Hollfelder at the University of Cambridge for valuable discussions and insights.

## Author contributions

**Kartik Totlani** : Writing - review & editing, Writing - original draft, Validation, Methodology, Investigation, Formal analysis. **Essa Ahsan khan** : Writing - review & editing, Writing - original draft, Validation, Methodology, Investigation. **Husnain Ahmed** : Writing - review & editing, Validation, Methodology, Investigation. **Rahmi Lale** : Writing - review & editing, Validation, Supervision, Resources, Funding acquisition, Conceptualization. **Bjørn T. Stokke** : Writing - review & editing, Validation, Supervision, Resources, Funding acquisition, Conceptualization.

## Electronic Supplementary Information

Electronic Supplementary Information (ESI†) is available, including detailed device schematics, coating protocols, additional microscopy images, and droplet stability experiments.

## Data Availability Statement

The data that support the findings of this study are available from the corresponding authors upon reasonable request.

## Notes

### Competing Interest Statement

The authors have declared no competing interest.

### Summary of Updates

Updated abstract, Introduction, revised Figure.3 and ESI files added

