## Supplementary material for "Integrated microdroplet workflow for high-throughput cell-free transcription in double emulsion picoreactors": Electronic Supplementary Information (ESI)

Supplementary Files :

1. A movie combining (i) plasmid compartmentalization, (ii) droplet re-injection, (iii) step-injection and (iv) double emulsification steps.
2. Stepinjection movies for the three design variations (A, B & C).
3. AutoCAD files for droplet formation and integrated reinjection-stepinjection and double emulsification devices.

| Primer Name | Sequence |
|---|---|
| Lib1a-F | CCTTTAATTAAATGTAAATGTTGCCATGTGTATGTG |
| Lib2a-R | TCCAAGACTAGTACGCGCTACATCCGCTTTAG |
| Lib1b-F | TAATACGACTCACTATAGGATGTAAATGTTGCCATGTGTATGTGG |
| Lib2b-R | ACGCGCTACATCCGCTTTAG |
| Lib3-F | GCCTTTAATTAATAATACGACTCACTATAGGATGTAAATGTTGC |
| Lib4-R | TCCAAGACTAGTACGCGCTACATCCGCTTTAGTATG |
| Lib5-F | GGATGTAGCGCGTACTAGTCTTGGACTCCTGTTGATAG |
| Lib6-R | GTGAGTCGTATTATTAATTAAAGGCATCAAATAAAACGAAAG |
| Lib7-F | GTCCTACTCAGGAGAGCGTTC |
| Lib8-R | GCCATGAATGATCCCGAAGG |

**Table S1** : List of primers used for plasmid construction and their respective sequences

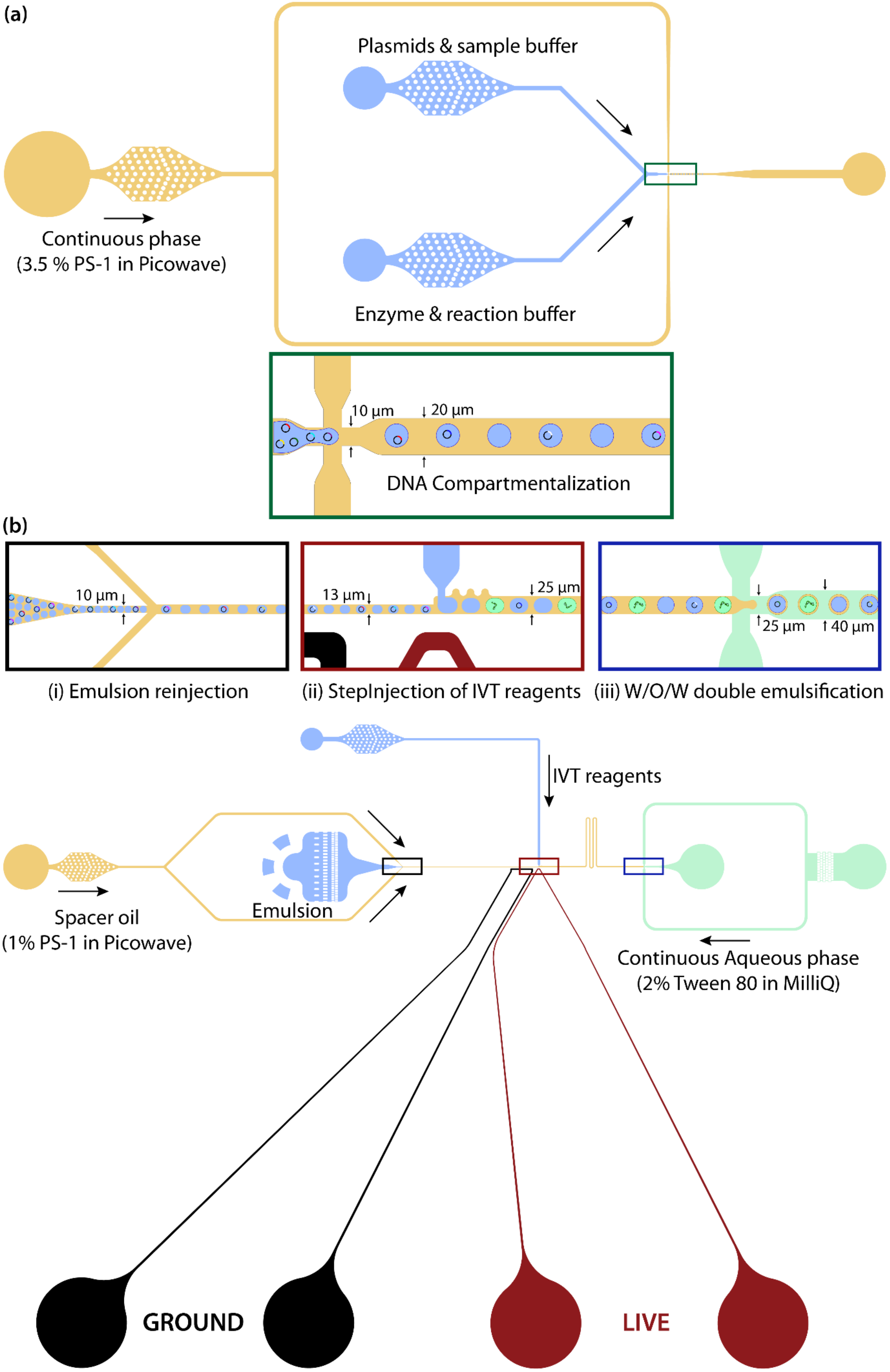
(a)
Plasmids & sample buffer
Continuous phase
(3.5 % PS-1 in Picowave)
Enzyme & reaction buffer
10 µm
20 µm
DNA Compartmentalization
(b)
10 µm
13 µm
25 µm
25 µm
40 µm
(i) Emulsion reinjection
(ii) StepInjection of IVT reagents
(iii) W/O/W double emulsification
IVT reagents
Spacer oil
(1% PS-1 in Picowave)
Emulsion
Continuous Aqueous phase
(2% Tween 80 in MilliQ)
GROUND
LIVE

**Figure.S1 : Design and layout of microfluidic devices.** (a) Microfluidic droplet generation device comprising of dual inlet for aqueous phases and single inlet for continuous oil phase. The key dimensions of the microfluidic channel are highlighted in the inset image (DNA compartmentalization). The height of the microchannel was 10.5 $\mu$m, while the nozzle is 10 $\mu$m wide and the main channel is 20 $\mu$m wide. (b) Integrated microfluidic device comprising of three key droplet operations (shown as insets): (i) Emulsion reinjection, (ii) IVT reagent stepinjection and (iii) Double emulsification. The dimensions of the various regions are stated in the insets. The microfluidic device is fabricated in two layers. Layer-01 which includes all the parts was 11 $\mu$m high and the second layer (which was exposed on the electrode regions and double emulsification part & inlets was 23 $\mu$m high (combined height of 34 $\mu$m). The re-injection nozzle is 10 $\mu$m wide the main channel is 13 $\mu$m wide before and 25 $\mu$m after the StepInjection junction. The red coloured channel indicates the live electrode whereas the black one indicates the ground electrode.

### Design variations for Stepinjection characterization

1. Design - A : $H_{main\ channel}$ = 8 $\mu$m; $W_{main\ channel-1}$ = 13 $\mu$m; $W_{main\ channel-2}$ = 23 $\mu$m; Nozzle = 10 $\mu$m.
2. Design - B : $H_{main\ channel}$ = 11 $\mu$m; $W_{main\ channel-1}$ = 15 $\mu$m; $W_{main\ channel-2}$ = 40 $\mu$m; Nozzle = 15 $\mu$m.
3. Design - C : $H_{main\ channel}$ = 11 $\mu$m; $W_{main\ channel-1}$ = 13 $\mu$m; $W_{main\ channel-2}$ = 25 $\mu$m; Nozzle = 10 $\mu$m.

### Microelectrode fabrication and device coating

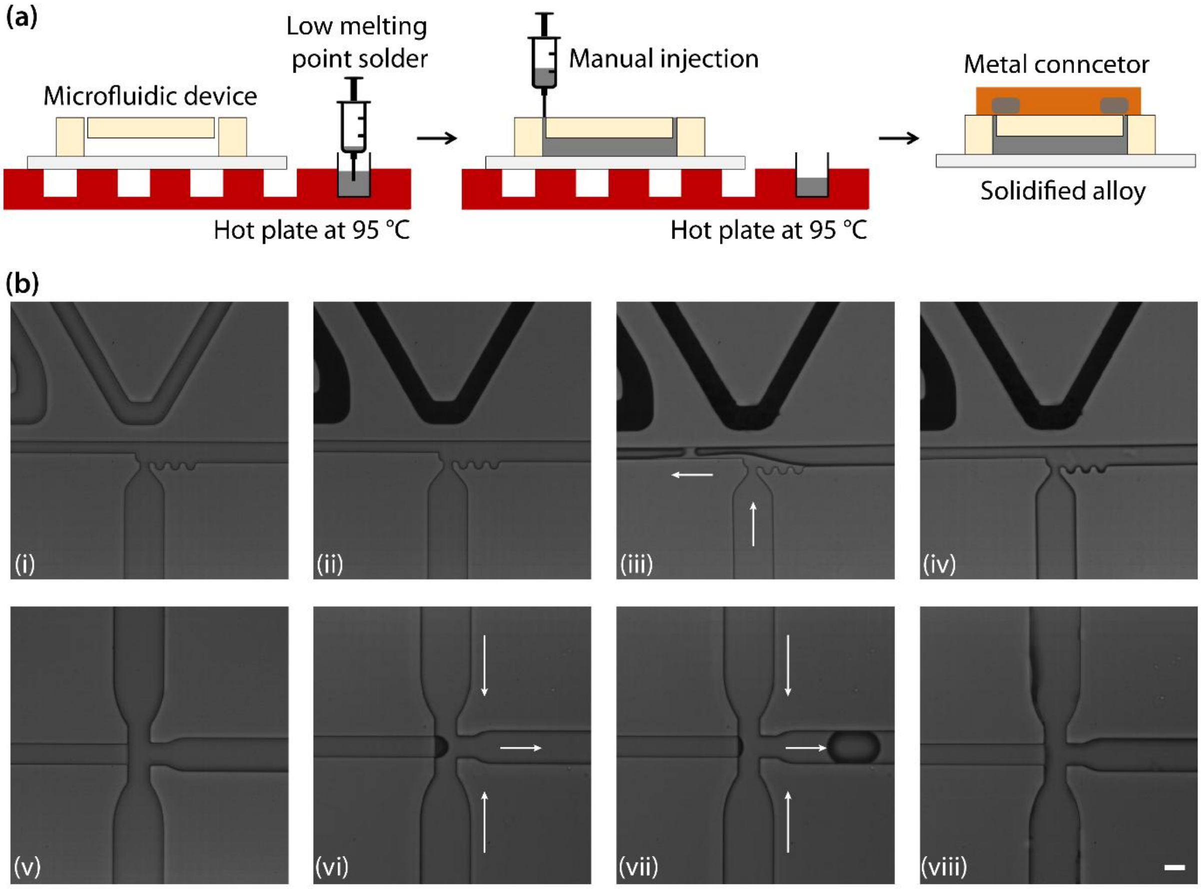


**Figure.S2 : Microelectrode fabrication and spatially selective surface modification of integrated step-injection devices.** (a) Schematic representation of the micro-electrode fabrication process. The microfluidic chip and molten alloy (Indalloy # 19 - 51In, 32Bi, 16.5Sn) are maintained at 95°C on a heater. The molten solder is injected into the designated electrode channels using a syringe until it emerges out from the outlet, ensuring complete and uniform filling. The electrode inlets and outlets are terminated with connector pins, and the device is removed from the heater for re-solidification. (b) Sequential grayscale images depicting the spatially selective surface modification process for integrated step-injection devices. **Top panel (i–iv):** Selective hydrophobic modification of the reinjection and StepInjection (SI) region using 1% (v/v) Perfluorooctyl-trichlorosilane in HFE-7500. (i) Bonded PDMS device prior to functionalization or electrode fabrication. (ii) PDMS device post microelectrode fabrication. (iii) Introduction of the silane into the SI junction; the immiscible oil-air interface advances through the channel under applied suction (white arrows), ensuring localized reagent confinement. (iv) Hydrophobic coating confined to the SI region, ensuring no cross-contamination to adjacent areas. **Bottom panel (v–viii):** Localized hydrophilic coating of the downstream flow-focusing (FF) junction via layer-by-layer (LBL) polyelectrolyte assembly. (v) Uncoated downstream channel prior to hydrophilic treatment. (vi) Controlled injection of PDADMAC/PSS polyelectrolyte solution with visible air–aqueous interface maintained to preserve channel wettability. (vii) Transient air bubble formation during withdrawal indicating active flow (white arrows). (viii) Finally, a hydrophilic (LBL coated) downstream FF junction for W/O/W emulsification. This selective patterning enables hydrophobic reagent injection and hydrophilic droplet pinch-off within the same device. Scale bar is 25 µm.

#### Droplet PCR Experiments with Q5 and TAQ

To assess the stability of surfactant-stabilized emulsions during thermocycling, droplet PCR experiments were performed using both Q5 (New England Biolabs) and Taq (Thermo Fisher Scientific) polymerase systems. The aqueous PCR mixtures were prepared with a total reaction volume of 200 µL for each formulation. For Q5 PCR, the mix contained 7.5 µL each of forward and reverse primers (Lib7 and Lib12), 10 µL of T7 Mango DNA template, 100 µL of Q5 master mix, and 75 µL of nuclease-free water. The Taq-based system consisted of 7.5 µL each of forward and reverse primers, 6 µL of DNA template, 20 µL of 10× buffer, 4 µL of dNTPs, 1 µL of Taq polymerase, and 154 µL of nuclease-free water.

Each premixed aqueous phase, containing the T7-Mango plasmid as template, was emulsified into picoliter droplets using a single-inlet flow-focusing microfluidic device (nozzle = 10 µm, main channel = 20 µm) with the continuous phase comprising either 5% PicoSurf-1 (Spherefluidics) or 4% FSO (Emulseo) dissolved in HFE-7500 oil. The resulting monodisperse emulsions were collected off-chip and subjected to thermocycling. For both systems, thermal cycling involved an initial denaturation at variable maximum temperatures (90 °C, 95 °C, or 98 °C) for 2 min, followed by 35 cycles of denaturation (10 s), annealing at 68 °C (20 s), and extension at 72 °C (60 s for Q5, 68 °C for Taq). The reactions were finalized with a 2–5 min hold at 72 °C and maintained at 4 °C thereafter.

To evaluate droplet stability, emulsions were imaged immediately after formation and again following completion of the PCR cycles using brightfield microscopy. Across both the tested surfactant formulations, droplet coalescence increased markedly with higher maximum denaturation temperatures, particularly above 90 °C (see figure S3 (a & b)). These observations qualitatively confirmed that thermal cycling compromises emulsion stability regardless of polymerase formulation, underscoring the advantage of isothermal amplification for preserving droplet integrity during DNA amplification.

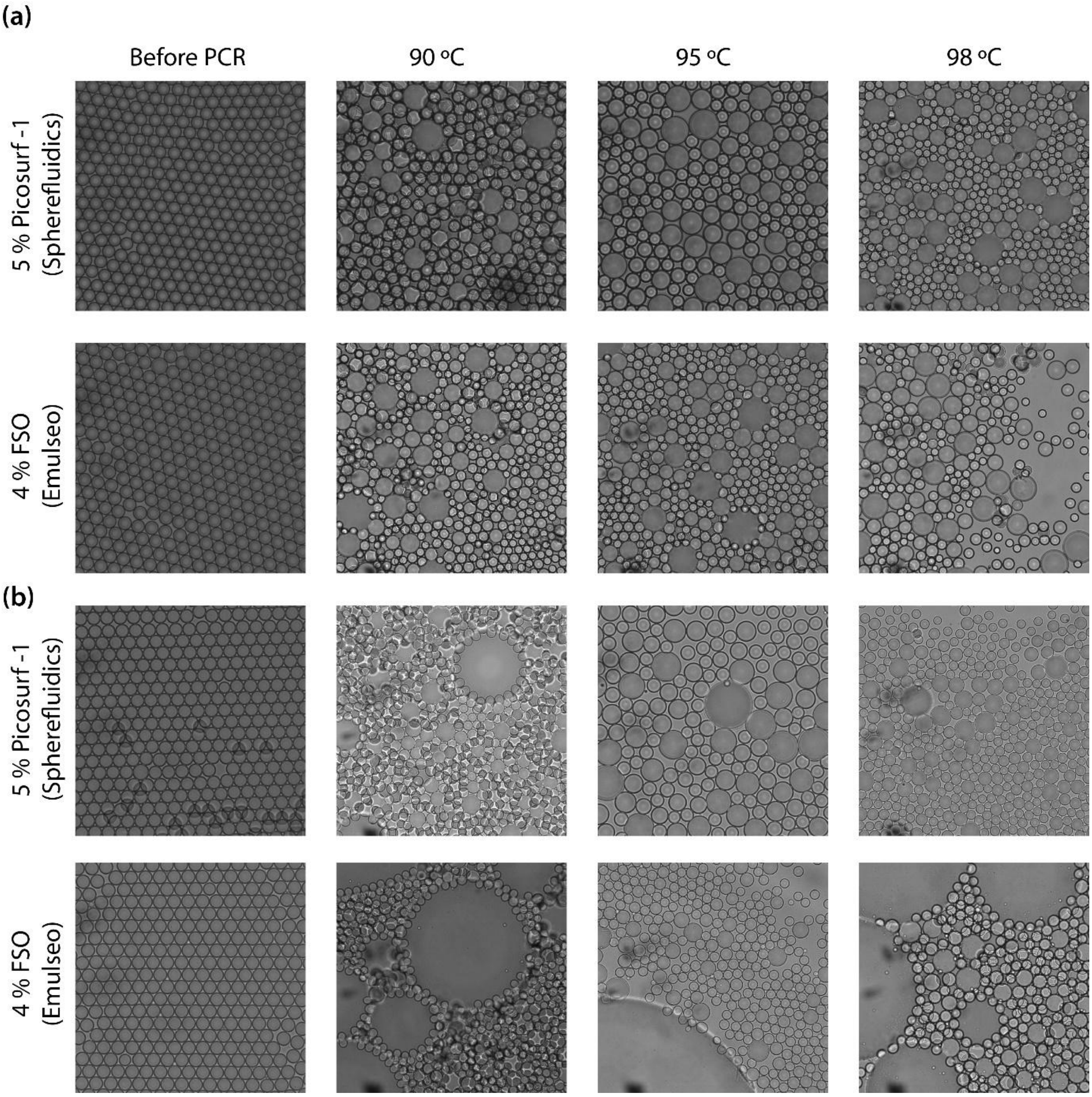


**Figure.S3 : Assessment of droplet stability during thermocycling under different temperatures and surfactant formulations.** (a) Microscopic greyscale images showing picodroplets emulsion before and after PCR thermocycling for droplets containing Q5 polymerase, encapsulated using two different oil–surfactant systems: 5 % PicoSurf-1 in HFE-7500 (top row) and 4 % Fluoro Surfactant (FSO) in HFE-7500 (bottom row). (b) Similar experiments using TAQ polymerase under identical conditions. In all cases, droplets were generated using a standard flow-focusing device producing ~ 2.5 pL droplets. Images in the left column show highly monodisperse, stable emulsions prior to thermocycling. Following thermal cycling (35 cycles, max temp : 90°C, 95°C, 98°C), noticeable coalescence, loss of monodispersity, and phase instability are observed varying in extent based on the surfactant and polymerase used.
